# NaVis: a virtual microscopy framework for interactive histological interrogation of spatial transcriptomics data

**DOI:** 10.64898/2026.02.18.706509

**Authors:** Ayomide Oshinjo, Jiahui Wu, Petar Petrov, Ali Hashmi, Johanna Englund, Valerio Izzi

## Abstract

Despite the widespread adoption of spatial transcriptomics (ST), revealing the alignment between transcriptional layers and tissue morphology remains technically demanding, typically requiring proficiency across multiple computational frameworks and thereby limiting accessibility for a substantial fraction of the biomedical community. Here, we introduce NaVis (https://github.com/Izzilab/NaVis), a point-and-click virtual microscopy framework that redefines ST analysis as an interactive, image-centric experience. NaVis enables rapid high-resolution inference from low-resolution whole-transcriptome platforms, producing microscopy-like visualizations while preserving transcriptome-wide coverage. It further decomposes histological images into quantitative tissue architecture priors – nuclei-rich regions, fibrillar extracellular matrix, and soft tissue – allowing direct integration of gene expression with local morphology. This unified representation supports analyses of compartment enrichment, boundary concordance, spatial cross-correlation, morphological patterning, histology–expression decoupling, and transcriptome-wide spatial similarity. By coupling transcriptomic and image-derived information within an interactive framework, NaVis shifts ST from static computational workflows to an exploratory modality, broadening its accessibility, conceptual reach and potential for biological discoveries.

---

Sequencing-based spatial transcriptomics platforms are rapidly becoming the method of choice for unbiased biological discovery^1^, further extending into clinical settings through FFPE-compatible protocols^2,3^. Platforms such as the 10x Genomics Visium family generate two intrinsically linked outputs from the same tissue section: spatially resolved gene expression profiles and a co-registered high-resolution histological image (typically hematoxylin and eosin, H&E). These modalities constitute complementary readouts of a shared biological substrate rather than alternative representations. Yet, most existing analytical frameworks either neglect the biological information encoded in the histological image, use it for visual overlays, or leverage it to refine operations on the transcriptomic layer, such as clustering, denoising, or dimensionality reduction^4–8^. In contrast, the reciprocal perspective – interrogating gene expression explicitly within its native histological context – remains markedly underexplored and largely confined to researchers with formal bioinformatics training^9^, limiting accessibility for pathologists, clinicians, and experimental biologists whose domain expertise is essential for biological interpretation.

Here, we introduce NaVis (https://github.com/Izzilab/NaVis), an online and locally deployable Shiny application that transforms spatial transcriptomics into an interactive, microscope-like experience. Upon launch, NaVis presents a full-screen canvas with microscope-inspired floating panels accessible via a navigation bar **(Figure 1)**.

**Figure 1.**
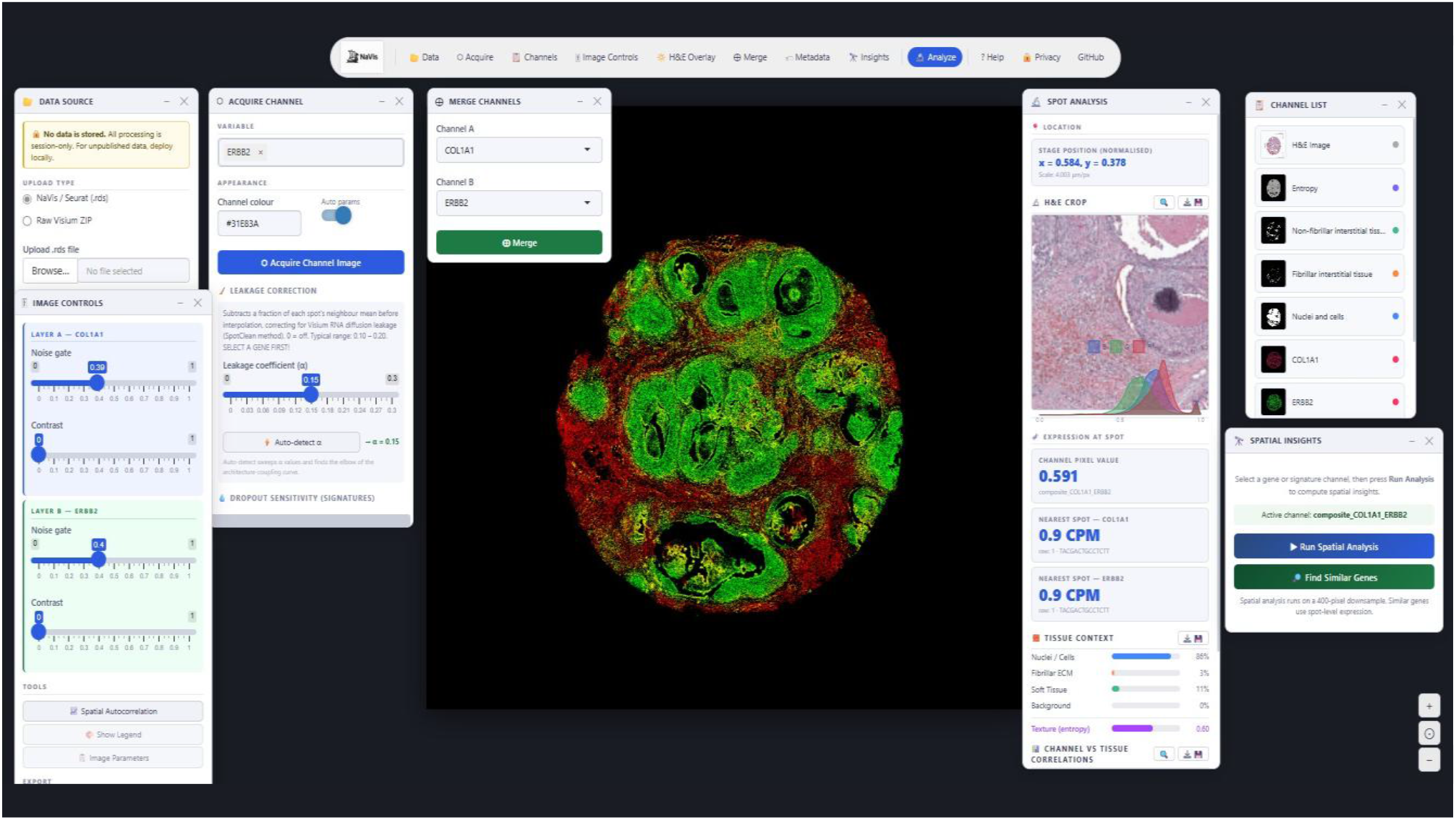
The NaVis interface.

The user interface is designed to minimize visual clutter and operate with robust default settings, while retaining full access to all algorithmic parameters **(Supplementary Methods)** through dedicated controls distributed across the panels. When a Visium dataset is loaded – either as a Seurat or SpatialExperiment object, a Space Ranger output folder, or via the bundled demo – the application automatically decomposes the H&E image into three quantitative tissue architecture compartments using color deconvolution, structure tensor, and texture analysis: a nuclei-rich compartment capturing cellular regions, a fibrillar compartment reflecting organized collagenous stroma, and a non-fibrillar compartment encompassing the remaining tissue. These components are exposed as independent channels within the interface *Channels* panel, alongside the original H&E image and any user-defined gene expression channels. Next, within the *Acquire* panel, users specify one or more genes, assign a display color (analogous to selecting a fluorescent probe in a microscope) and initiate channel generation via *Acquire Channel Image*. NaVis reconstructs continuous expression maps from the discrete spatial capture locations using an adaptive-decay inverse distance weighting (AD-EBIDW) scheme, further refined through image prior–informed spatial ridge regression **(Supplementary Fig. 1)**. Additional controls to mitigate RNA diffusion between adjacent spots using the SpotClean method^10^ and sensitivity to dropout events are available within the same panel, alongside full access to AD-EBIDW parameters **(Supplementary Fig. 2)**. The *Image Controls* panel provides per-channel contrast adjustment and noise gating, and dynamically expands to multi-channel controls when channels are combined through the *Merge* panel. This enables dual-channel compositing in complementary colors, recapitulating the appearance of fluorescence co-staining while also generating an agreement map (white for concordant signal, dark for discordant regions) and an interface map of where signals are juxtaposed but not overlapping (**Supplementary Fig. 3)**. Similarly, the *H&E Overlay* panel supports direct blending of gene and/or composite channels onto the histological image with real-time opacity and brightness control, while the *Metadata* panel further allows rendering of native-resolution spatial maps for both numerical and categorical annotations embedded in the data object.

Beyond a microscope-inspired discovery workflow, the central innovation of NaVis lies in its analytical framework which explicitly embeds gene expression within its native biological context, enabling the systematic interrogation of transcriptional patterns in direct relation to underlying tissue architecture. Pressing *Analyze* turns the cursor into a crosshair, and a click on the tissue opens a drawer that reports an H&E crop centered on the location, the RGB color histogram of the local histology, the percentage tissue composition (summing over the three compartments to 100%), the CPM-normalized expression of every gene in the active channel, and the Pearson correlation of the active channel image with each tissue prior. The Analyze environment is context-aware: a left-click on a channel shows these values, while a right-click stores the measurements in memory for comparison with a second set of measures from another area. Finally, shift-dragging the cursor opens the region-of-interest (ROI) panel, which provides the H&E crop of the ROI, percentage tissue composition, and the top-20 enriched genes (inside ROI *vs*. rest of the tissue) **(Supplementary Fig. 4)**.

The *Insights* panel extends the analytical framework by quantifying the relationship between gene (or composite) maps and tissue architecture across five complementary measures: (i) compartment enrichment, expressed as a fold-ratio with permutation-based significance; (ii) expression boundary alignment, capturing the degree to which transcriptional patterns conform to or traverse histological borders; (iii) spatial cross-correlation across increasing lag distances (0–800 µm), enabling scale-resolved association profiling; (iv) expression island morphology, characterizing connected high-expression regions (≥75th percentile) in terms of area, abundance, and circularity; and (v) histology decoupling, quantifying the extent to which spatial gene expression patterns diverge from those predicted by underlying histological structure. From the same panel, users can launch a BLAS-accelerated transcriptome-wide search for the ten genes whose spot-level expression profile most closely matches the query by pressing the *Find Similar Genes* button (**Supplementary Fig. 5)**.

To assess whether NaVis, although not designed as a super-resolution method, produces gene expression maps that faithfully recapitulate true sub-spot spatial structure, we benchmarked its output against single-cell Xenium in situ sequencing data from a matched human FFPE breast cancer section (10x Genomics; n = 167,780 cells, 313-gene panel)^11^. Visium spot expression was super-resolved with NaVis defaults and registered to the Xenium frame; 306 panel genes were shared between platforms. NaVis maps showed substantially stronger agreement with Xenium ground truth than either of two null baselines: median per-gene Pearson r = +0.246 against Xenium, versus +0.029 for raw Visium spot counts rasterized without smoothing and +0.052 for a gene-label–shuffled negative control **(Supplementary Fig. 6A)**. NaVis outperformed the naive baseline in 291 of 306 genes (95%; median Δr = +0.21; Wilcoxon signed-rank p = 1.1 × 10^−49^; **Supplementary Fig. 6B**), confirming that the gain reflects the super-resolution model itself rather than the underlying spot grid. Structural similarity (SSIM = 0.43 ± 0.08, mean ± s.d.; 70% of genes > 0.4; **Supplementary Fig. 6C**) and top-region concordance (median IoU of top-20% pixels = 0.19, 1.7× the chance level of 0.11; 41% of genes > 2× chance; **Supplementary Fig. 6D**) provided two metric-independent validations of the same conclusion, and SSIM correlated with Pearson r per gene (r = +0.30, p = 1 × 10^−7^; **Supplementary Fig. 6E**), arguing against artefacts of any single metric. Cross-platform pseudobulk panel-wide expression was strongly concordant (Pearson r = 0.853 on log-transformed totals; Fig. 5), establishing baseline agreement on gene abundances before spatial comparison. The magnitude of NaVis’s gain over the naive baseline scaled with Xenium transcript abundance (r = +0.43, p = 4 × 10^−15^; **Supplementary Fig. 6F**), indicating that NaVis recovers most additional information for genes in the regime where ground truth itself is well-sampled. Together, these results demonstrate that NaVis output is structurally faithful to single-molecule ground truth despite not being optimized for spatial reconstruction, and that the limits of its agreement with Xenium are determined by Visium’s intrinsic resolution rather than by the super-resolution step. Importantly, existing approaches for enhancing Visium data typically do not operate directly on native Seurat objects with embedded low-resolution images. In contrast, NaVis maintains compatibility with these standard data structures and enables visual resolution enhancement by leveraging histology-guided decomposition **(Supplementary Fig. 7)**. This approach allows users to generate higher-resolution visualizations from low-resolution data, thereby facilitating seamless integration with the large body of datasets already available in public repositories.

In summary, NaVis reframes ST as a microscopy-like, image-centric experience that bridges transcriptomics and computational pathology while lowering computational barriers. By enabling immediate, interactive access to spatial inference, it broadens ST usability across clinicians, pathologists, experimental biologists, and computational researchers, and extends both the practical applicability and conceptual scope of spatial omics.

## Author contributions Statement

V.I. conceived the study and developed the conceptual framework underlying NaVis. A.O., J.W., P.P. and V.I. designed the computational architecture. A.O., A.H. and V.I. performed analyses and developed the GitHub repository. V.I. and J.E. wrote the manuscript with input from all authors. All authors reviewed, edited, and approved the final manuscript.

## Data availability

All datasets used for demonstration are publicly available from 10x Genomics (https://www.10xgenomics.com/datasets/) or selected publications (https://www.nature.com/articles/s41467-023-43458-x). Further preprocessed datasets in NaVis format are accessible from Figshare (https://figshare.com/s/3e568dcb32106767cd5e) [DOI: 10.6084/m9.figshare.31111300].

## Code availability

The NaVis application is freely accessible online at https://matrinet.shinyapps.io/NaVis. Local installation package and detailed documentation are available at https://github.com/Izzilab/NaVis.

