## Supplementary Figures for "NaVis: a virtual microscopy framework for interactive histological interrogation of spatial transcriptomics data"

[+]equally-contributing authors

### Supplementary Figures

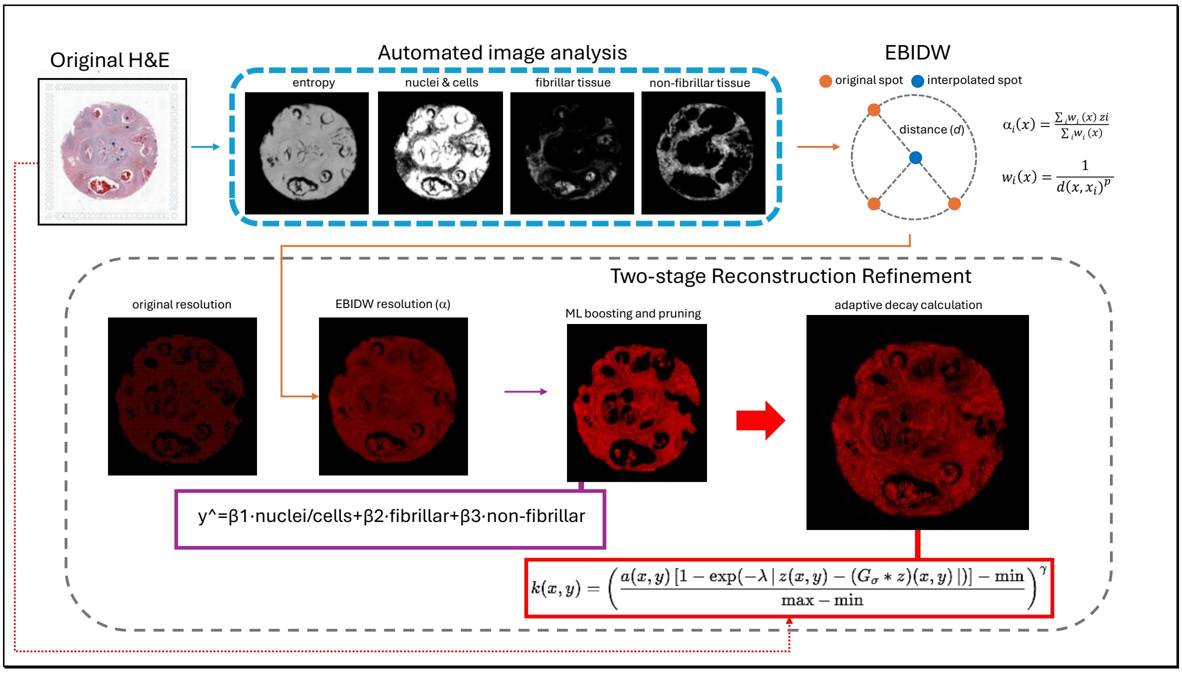


**Supplementary Figure 1. Overview of the NaVis algorithm.**
After data upload, NaVis initiates its integrated image-analysis modules, which are optimized for rapid extraction of nuclei/cell features and fibrillar and non-fibrillar tissue components. Expression-based inverse distance weighting (EBIDW) is first applied to generate a smoothed interpolation ($\alpha$) of the original coarse-resolution gene expression map. This interpolated image is then passed to a ridge regression-like machine learning module with L2 regularization, which decomposes the signal into its underlying biological components and adaptively boosts or prunes each according to its inferred contribution, while incorporating an entropy estimate to capture local uncertainty. The resulting image is finally refined using a decay mask—computed as the difference between a reference image (the H&E image by default) and its Gaussian-blurred counterpart—to selectively suppress interpolation and ML-derived artefacts and produce the final high-resolution rendering. All EBIDW parameters, along with the option to enable the ML model or to select an alternative decay mask, can be fully adjusted by the user.

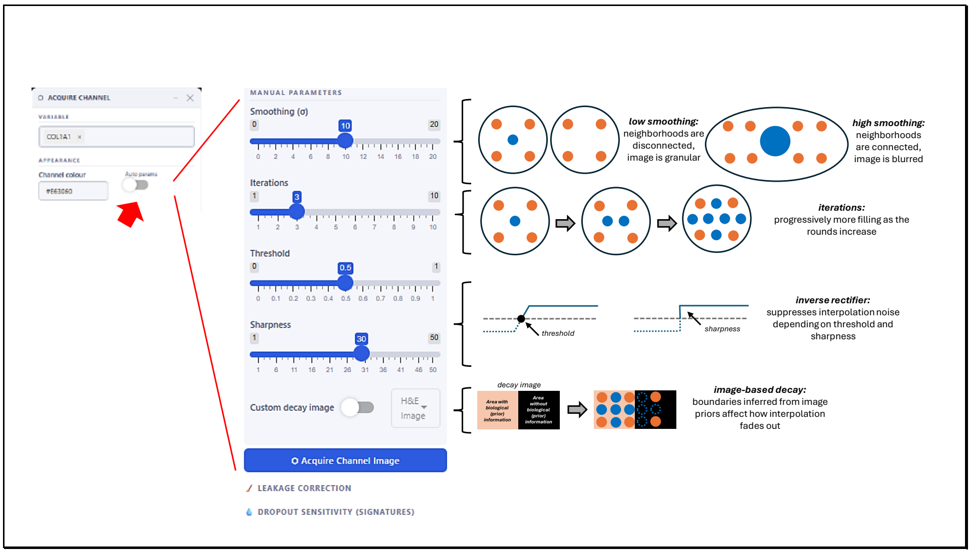


**Supplementary Figure 2. AD-EBIDW parameters.**
AD-EBIDW parameters are not available by default, but users can access them by clicking the ”Auto params” button in the ”Acquire Channel” panel. Smoothing, number of iterations, and the threshold and sharpness of the noise-suppressing rectifier stage can be controlled. Also, users can choose a different decay image to be used for the final stage of the algorithm. The default parameters used by NaVis are shown. Note that the controls for “Leakage correction” using the SpotClean method and “Dropout sensitivity” (for signatures) are always accessible and shown to the user.

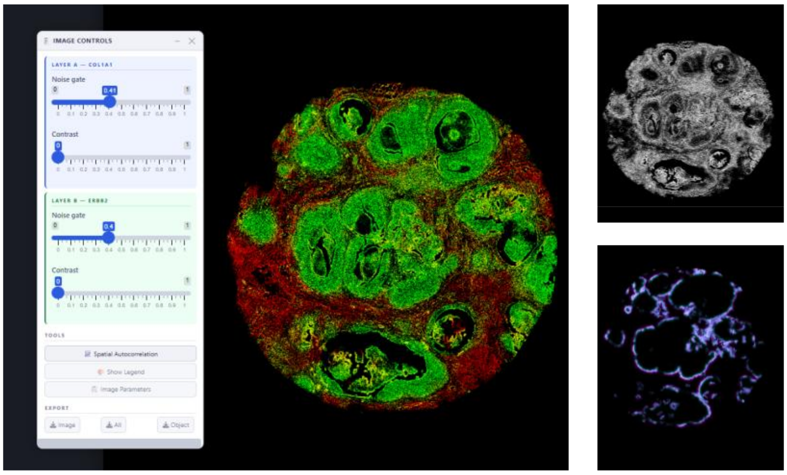


**Supplementary Figure 3. Image controls.**
A) The “Image controls” panel allows access to parameters for image appearance, such as noise gate and contrast, and changes in real time according to whether the image is a single channel ora composite of two channels (left, shown for a composite of *COL1A1* and *ERBB2*). When forming a composite, also B) a grayscale agreement map (black: no agreement, white: full agreement) and C) an interface map (areas where two channels face each other without overlapping) are generated.

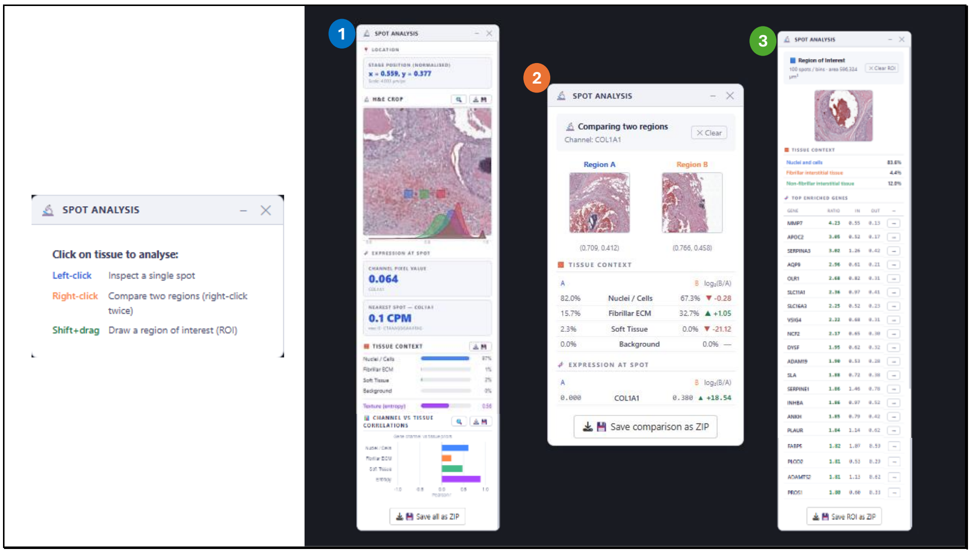


**Supplementary Figure 4. Analysis modes.**
The “Analyze” panel is context-aware and allows for three different models of analysis. A single, left-click on any point of the image (1, blue) opens a panel with information on analysis spot location, H&E crop, RGB histogram, gene expression at location and correlation between gene expression and image priors. Two right-clicks (2, orange) opens a comparison tab, with H&E crops, relative enrichment or depletion of image priors, and comparative expression. Finally, shift-dragging draws a region of interest (ROI) showing a H&E crop with dimensions, tissue context, and top 20 most-enriched genes (inside the ROI *vs.* outside).

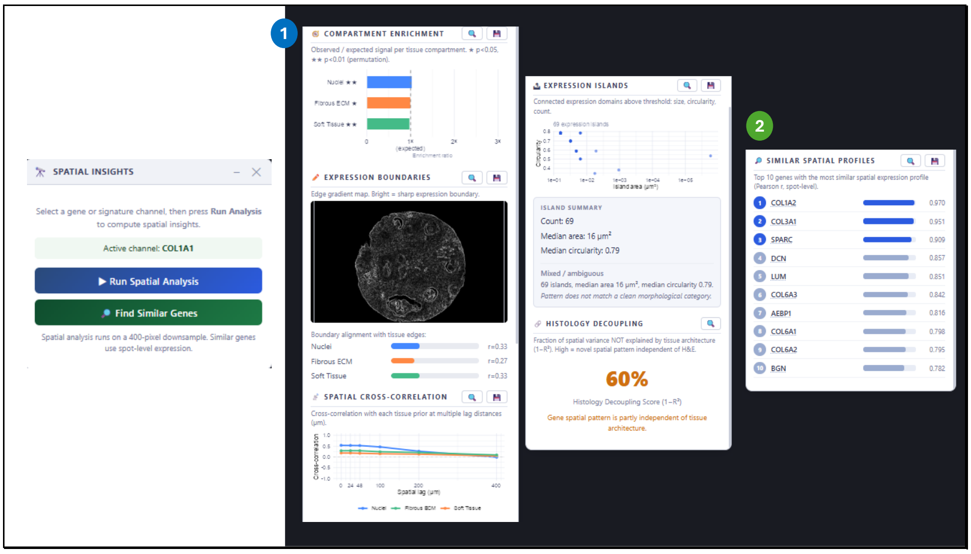


**Supplementary Figure 5. Insights.**
The "Insights" panel allows image-wide analyses of the relationship between gene expression maps and the H&E histology map. The "Run Spatial Analysis" button (1, blue) evaluates gene expression enrichment *vs.* image prior using a permutation-based test, boundary analysis (tissue edges highly correlating with gene expression), spatial cross-correlation at lags, expression islands and histology decoupling. The "Find Similar Genes" button (2, green) performs image-wide Pearson correlation tests for gene expression across .

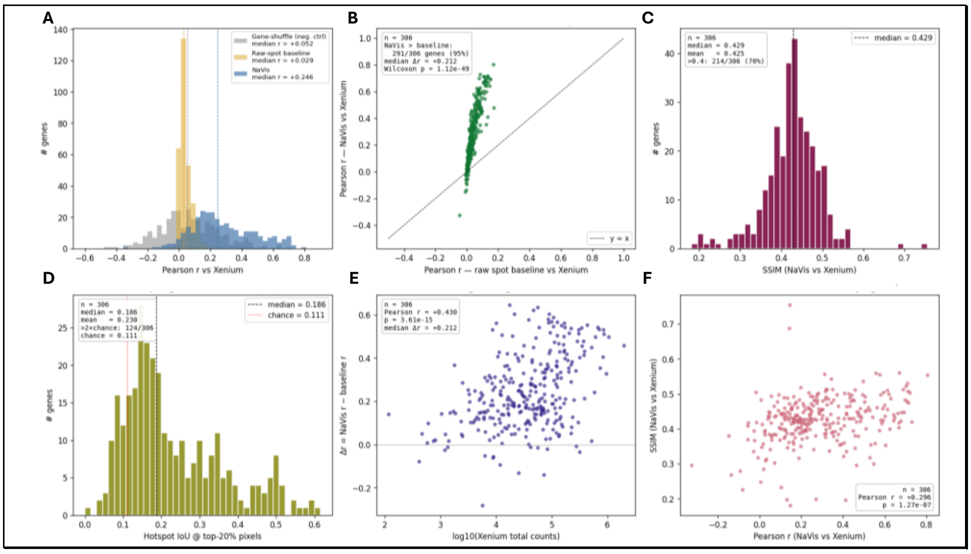


**Supplementary Figure 6. NaVis comparison vs. matching Xenium.**
Comparison metrics for a matching Xenium-Visium dataset (GSE243280).

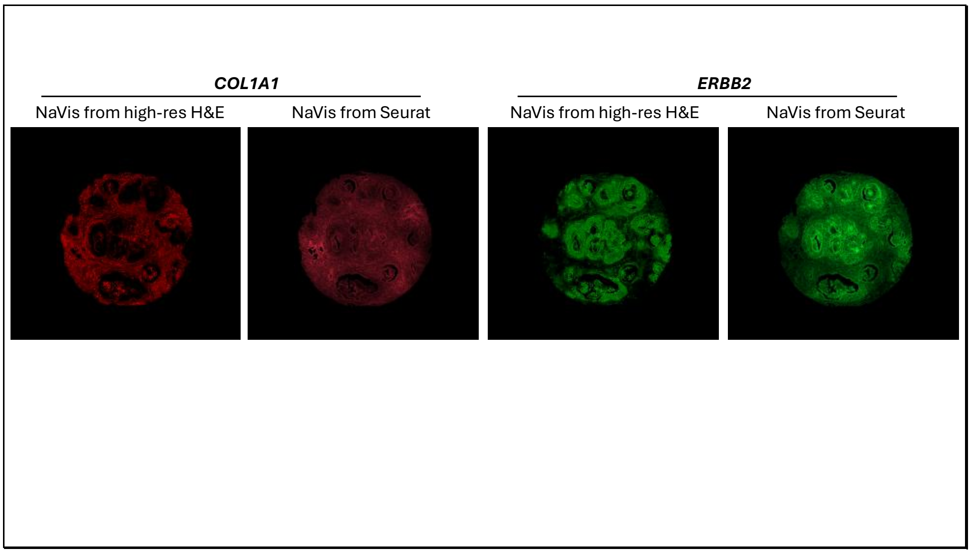


**Supplementary Figure 7. Comparison of NaVis visualizations from high- and low-resolution Visium images.**
Unlike other methods focusing on increasing the resolution of Visium datasets, NaVis can operate directly on standard Seurat and SpatialExperiment objects, typically built from low-resolution H&E images.
