## Supplementary Methods for "NaVis: a virtual microscopy framework for interactive histological interrogation of spatial transcriptomics data"

[+]equally-contributing authors

### Supplementary Methods

#### S1. Overview of the pipeline

NaVis converts a sequencing-based spatial transcriptomics (ST) experiment, most typically a 10x Genomics Visium dataset, into a coherent set of two-dimensional image channels defined on a common pixel grid, and makes them interactively analyzable through a panel-based interface. This Supplementary Methods document describes every transformation the tool applies, in the order in which they run when data is loaded and a user issues commands. Each section contains: (i) the mathematical definition of the operation, (ii) the parameter choices and their justification, (iii) the implementation complexity, and (iv) edge cases and failure modes.

Throughout, we denote the Visium expression matrix as $X \in \mathbb{R}^{G\times N}$, with $G$ genes indexed by $g$ and $N$ spots indexed by $i$. The spatial coordinates of spot $i$ in the H&E image plane are $\mathbf{c}_i = (x_i, y_i)$. The H&E image itself is a three-channel RGB array $H \in [0, 1]^{H\times W\times 3}$. All image reconstructions produced by NaVis are also defined on the same $H\times W$ grid.

### S2. Data ingestion and normalization

NaVis accepts three input formats: a Seurat object (v4 or v5) saved as RDS, a SpatialExperiment object saved as RDS, or a Space Ranger output directory (including the CytAssist variant with parquet-format "tissue_positions.parquet" spatial locations). Further platforms that can be represented as a Seurat object are technically compatible with NaVis, as long as a histological image (low- or high-res) is embedded within. Currently, *in situ*-based methods that do not include an image in Seurat format (such as Xenium) are not supported. From the uploaded material, NaVis extracts four components: the sparse expression matrix $X$, the barcode-indexed spot coordinates, the H&E hi-res or low-res image, and the Visium scale factor sf that converts pixel-space coordinates to their actual resolution equivalent.

#### S2.1 Automatic normalization fallback

When Space Ranger output directory is used as input, NaVis perfroms all the regula steps to build a Seurat object, including normalization. When the uploaded object is a Seurat file that has not been normalized, the `data` layer is empty; in that case NaVis reads the raw `counts` layer and applies the same log-CPM normalization that Seurat applies by default:

$$\tilde{X}_{g,i} = \log(1 + 10^4 \cdot X_{g,i} / (\sum_g X_{g,i}))$$

For Seurat v5 objects, layers are accessed through the object's *`Layers`* slot and do not carry rownames; in this case, NaVis reattaches feature names and spot barcodes from the assay's own rownames before returning the normalized matrix. Objects that, for any reason, pass an unnamed matrix to downstream operations return an error message and are not allowed to continue.

### S3. H&E decomposition into tissue-architecture priors

NaVis decomposes the H&E image into three compartment maps $M_{nuc}$, $M_{fib}$ and $M_{soft}$ plus an entropy texture map $M_{ent}$. The first three are the quantitative priors used by the *Insights* readouts; the entropy map is reported separately as a texture feature.

#### S3.1 Colour deconvolution for nuclei and eosin

We separate H&E staining into hematoxylin (nuclei) and eosin (cytoplasm / stromal protein) channels via Beer–Lambert color deconvolution with the standard stain matrix of Ruifrok and Johnston (Ruifrok & Johnston 2001).

$$O = -\log(H + \varepsilon), \quad S = (M_{RJ})^{-1} O$$

where $M_{RJ}$ is the 3×3 Ruifrok–Johnston stain matrix whose rows are the normalized RGB absorbance vectors of hematoxylin, eosin and the null channel, and $\varepsilon = 10^{-6}$ prevents log(0). The two stain channels $S_H$ and $S_E$ are re-normalized to [0, 1].

The nuclei compartment is built from $S_H$ by Otsu thresholding followed by morphological opening and hole-filling to remove salt-noise and fill single-pixel gaps inside nuclei:

$$M_{nuc} = \text{fill\_hull}(\text{open}(S_H \geq \tau_{Otsu}, \text{brush\_5}))$$

followed by connected-component labelling (`EBImage::bwlabel`). Each nucleus receives a unique integer label. The label image is what is stored in $M_{nuc}$; wherever nuclei density is needed the image is binarised ($M_{nuc} > 0$), and wherever per-nucleus identity is needed the labels are used directly.

### S3.2 Fibrillar ECM from structure tensor and Gabor filters

Fibrillar extracellular matrix appears in H&E as organized, directionally coherent eosinophilic structure. We quantify this with a weighted combination of three complementary features: structure-tensor coherence, directional Gabor-filter response, and eosin density.

#### *Structure-tensor coherence*

For each pixel, we compute the smoothed 2×2 structure tensor $J$ from the grayscale gradient:

$$J(p) = G_\sigma * ((\nabla I)(\nabla I)^T)(p)$$

with Gaussian smoothing kernel $G_\sigma$ of radius $\sigma = 2$ pixels. Let $\lambda_1 \geq \lambda_2$ be the eigenvalues of $J$. The coherence measure is:

$$\text{coh}(p) = ((\lambda_1 - \lambda_2) / (\lambda_1 + \lambda_2 + \varepsilon))^2$$

with $\varepsilon = 10^{-8}$, coherence is close to 1 where the local gradient is strongly oriented (collagen fibers) and close to 0 in isotropic or uniform regions.

#### *Gabor-filter energy*

A Gabor filter bank samples oriented, band-limited texture at multiple scales. We use $S = \{2, 4\}$ pixel wavelengths and $\Theta = \{0, 45°, 90°, 135°\}$ orientations. The total Gabor energy is the sum of squared responses of the real-part filters at each (scale, orientation) pair, normalized by the maximum over the image:

$$\text{gab}(p) = \sum_{s \in S} \sum_{\theta \in \Theta} (g_{s,\theta} * I)(p)^2$$

followed by $L\infty$-normalization over the image.

#### *Combined fibrillar ECM score*

The three components are combined by convex combination with weights chosen empirically from a range of breast, colon and brain sections:

$$M_{fib} = 0.5 \cdot \text{coh} + 0.3 \cdot \text{gab} + 0.2 \cdot S_E$$

The weights 0.5 / 0.3 / 0.2 were selected so that (a) coherent oriented structure dominates the signal (coherence is the most specific feature of organized stroma), (b) Gabor energy captures mid-scale texture missed by the eigenvalue ratio, and (c) eosin density breaks ties between collagen-poor soft tissue regions and stroma-poor epithelial regions. The combined map is returned in [0, 1] after global min-max normalization.

#### S3.3 Soft tissue by residual diffusion

Soft tissue is defined as the complement of nuclei and fibrillar ECM, computed as a smooth residual via Laplace diffusion from nuclei seeds modulated by the fibrillar map. Let $u$ be the solution of the inhomogeneous Laplace equation:

$$(1 - \alpha M_{\text{fib}}) \cdot \Delta u = M_{\text{nuc}}, \quad u = 0 \text{ on } \partial\Omega$$

with $\alpha = 0.7$ set by the proportionality of fibrillar structure needed to keep soft tissue from leaking into stroma, solved by 50 iterations of the Gauss–Seidel scheme on the pixel grid. The solution $u$ is a smoothly varying field that is high in nuclei-rich regions and attenuated where fibrillar structure is strong; its complement after normalization is the soft-tissue density:

$$M_{\text{soft}} = 1 - \text{normalize}(u)$$

#### S3.4 Local entropy texture

The entropy map is Shannon entropy of the 8-bin grayscale histogram computed in a 9×9 sliding window:

$$M_{\text{ent}}(\text{p}) = -\sum_{k=1..8} p_k(\text{p}) \cdot \log_2 p_k(\text{p})$$

where $p_k(\text{p})$ is the fraction of pixels in the window centered on p that fall in the $k$-th intensity bin. High entropy flags textured regions (*e.g.* inflamed tissue, tumor–stroma interfaces) that are not reducible to a single compartment. The map is reported separately as a texture feature; it does not participate in the Insights readouts but is available in the click-to-inspect drawer.

### S4. Adaptive-decay EBIDW interpolation

Each gene's discrete location-level expression is lifted to a continuous image via an Adaptive-Decay Expression-Based Inverse Distance Weighting (AD-EBIDW) scheme. The goal is to produce a continuous spatial field that resembles a fluorescent stain, yet (i) agrees with the measured values at each spot, (ii) decays smoothly between spots at a rate matching the Visium resolution, and (iii) suppresses noise at low expression without dampening strong local peaks.

#### S4.1 Spot rasterization

For a gene $g$, spot values $X_{g,i}$ are projected onto the H&E pixel grid at each spot's integer coordinates:

$$S_g(\text{p}) = X_{g,i} \text{ if } \text{p} = \text{round}(c_i) \text{ for some } i, \quad \text{else } 0$$

This produces a sparse image of non-zero values at $N$ discrete positions on an $H \times W$ grid.

### S4.2 Iterated isotropic Gaussian diffusion

The sparse image is diffused by repeated application of an isotropic Gaussian kernel with base standard deviation $\sigma_0$ (default 7 pixels, ≈55 µm at Visium low-res scale factor) over $n_{\text{iter}}$ iterations (default 3). The result is equivalent to a single convolution with effective standard deviation

$$\sigma_{\text{eff}} = \sigma_0 \cdot \sqrt{n_{\text{iter}}}$$

because variances add under Gaussian convolution. Iterating rather than using a single larger kernel allows the per-iteration $\sigma_0$ to remain small, while the total diffusion distance matches the Visium spot-to-spot spacing. The smoothed image is thus:

$$\tilde{S}_g = (G_{\sigma_0})^{*n\_iter} * S_g$$

where $(G_{\sigma_0})^{*n\_iter}$ denotes $n_{\text{iter}}$-fold convolution of the Gaussian kernel with itself. In practice the iterated blur is computed directly as $n_{\text{iter}}$ successive calls to `imager::isoblur` for cache-friendly execution and progress reporting.

### S4.3 Sigmoid rectifier

Raw Gaussian smoothing attenuates low-expression noise uniformly, which produces a muddy image with low contrast between signal and background. NaVis applies a sigmoid rectifier that preserves peaks while sharpening the transition to near-zero values:

$$E_g = \tilde{S}_g \cdot (1 + \exp(s \cdot (t - \tilde{S}_g)))^{-1}$$

with default steepness $s = 10$ and threshold $t = 0.2$. The rectifier multiplies the smoothed image by a sigmoid gate centred at $t$, so that values well above threshold pass unchanged (gate ≈ 1) and values well below threshold are attenuated to near zero (gate ≈ 0). Unlike a hard threshold, the sigmoid preserves derivative information near the cutoff, which is important for the Sobel gradient readout (see S6.2).

Parameter $t$ is exposed to the user via the *Noise gate* slider in the Image Controls panel; $s$ is fixed. For signatures and in the "machine-learning correction" mode $t$ defaults to 0.2; for raw mode $t$ is user-controlled, default 0.3.

### S4.4 Complexity

The dominant cost of AD-EBIDW is the iterated Gaussian blur at $O(n_{\text{iter}} \cdot HW \cdot k^2)$ for a separable Gaussian kernel of support $k = \lceil 6\sigma_0 \rceil + 1 \approx 43$ pixels, implemented as two 1D passes for $O(n_{\text{iter}} \cdot HW\, k)$ total. For a typical 2000×1800 high-res H&E and defaults, one gene map takes ~2–3 seconds on a laptop CPU; signatures of $K$ genes cost $K\times$ that, serialized.

### S5. Spatial leakage correction

FFPE Visium sections exhibit RNA diffusion across spot boundaries, a phenomenon known as spot swapping ([Ni et al. 2022]). Each spot's observed expression is contaminated by a fraction of its neighbours' expression. We correct this directly on the spot-level matrix, before interpolation, using a SpotClean-inspired formulation.

#### S5.1 Model

Let W be a sparse $N \times N$ adjacency matrix where $W_{ij} = 1 / k$ if spot $j$ is among the $k = 6$ Euclidean-nearest spots of $i$, else 0. Row sums of W are 1 by construction, so W$X$ is the neighbor-averaged expression per spot. The corrected expression at strength $\alpha \in [0, 1]$ is

$$\hat{X}(\alpha) = \max(0, X - \alpha \cdot \mathrm{W}X)$$

Clipping to zero prevents negative values where local means exceed the spot value — these are biologically meaningless and would create phantom negative signal in the interpolation. $\alpha = 0$ disables correction; $\alpha = 1$ subtracts the full neighbor mean and is typically too aggressive (over-corrects in genuinely uniform regions).

#### S5.2 Auto-detecting α by Kneedle elbow

The optimal α depends on the specific gene and section. We auto-detect it by finding the point of diminishing return on a coupling-quality curve.

Sweep $\alpha$ over a small grid $A = \{0, 0.05, 0.10, 0.15, 0.20, 0.25\}$. For each α, compute a fast EBIDW map with reduced settings ($\sigma_0 = 5$, $n_{\mathrm{iter}} = 1$, 300×270 pixel downsample) and evaluate its coupling to tissue architecture by the adjusted $R^2$ of the linear regression

$$E_g(\alpha) \sim \beta_0 + \beta_1 M_{\mathrm{nuc}} + \beta_2 M_{\mathrm{fib}} + \beta_3 M_{\mathrm{soft}} + \varepsilon$$

on a 10%-subsample of pixels (uniform stride). This yields a curve $R^2(\alpha) \in [0, 1]$. As α increases from 0, leakage is removed and the gene/tissue architecture relationship tightens ($R^2$ rises). At some α*, further correction starts removing genuine signal and $R^2$ plateaus or decreases. We identify α* as the Kneedle elbow [Satopää et al. 2011] of the $R^2(\alpha)$ curve:

$$\alpha^* = \mathrm{argmax}_{\alpha \in A}\, d(\alpha)$$

where $d(\alpha)$ is the perpendicular distance from $(\alpha, R^2(\alpha))$ to the chord connecting the endpoints $(0, R^2(0))$ and $(\alpha_{\max}, R^2(\alpha_{\max}))$. This is the standard Kneedle criterion: the elbow is the point of maximum curvature on a monotone curve, and it marks the transition from meaningful correction to over-correction.

Computation time for auto-α is $O(|A| \cdot t_{\mathrm{EBIDW,fast}}) \approx 5$–8 seconds total on a laptop. The user invokes it via the *Auto-detect α* button; the recovered α* populates the slider and the Acquire button then runs at full settings with that α.

### S6. Dropout-aware co-expression gating for signatures

A co-expression signature is built from a set $G_{sig} = \{g_1, \ldots, g_K\}$ of $K \geq 2$ genes. Naively averaging the per-gene interpolated maps is inadequate for two independent reasons: (i) spots where one gene is zero would still contribute to the signature through neighbor diffusion, producing halos outside the true co-expression footprint; (ii) strict AND-gating ("any gene zero → spot excluded") eliminates genuine co-expression at spots where one signature gene has dropped out by chance, a dominant failure mode on Visium data.

NaVis addresses both concerns with a three-layer union-rule gating scheme.

#### S6.1 Layer 1 — dropout-aware true-zero classification

For each signature gene *g* and each spot *i*, classify the spot as true zero or dropout using 6-nearest-neighbour support. Let $N_6(i)$ be the 6 spatial nearest neighbors of *i*.

$$\text{is\_true\_zero}(i, g) = (X_{g,i} = 0) \wedge (|\{j \in N_6(i) : X_{g,j} = 0\}| / 6 \geq \tau)$$

where $\tau \in [0, 1]$ is the *Dropout sensitivity* slider (default 0.5). A spot is a true zero only if the gene is zero at the spot AND at least a τ fraction of its neighbors are also zero. Otherwise, the zero is classified as a dropout artefact and imputed as the mean of non-zero neighbor values:

$$X_{g,i} \leftarrow \text{mean}\{X_{g,j} : j \in N_6(i), X_{g,j} > 0\}$$

The imputed values are written back into the expression matrix (a local copy, not the user's original object).

#### S6.2 Union rule: reject if all genes are true zero

Let $T_g$ be the set of spots classified as true zeros for gene *g*. Define the signature's non-coexpression set as the intersection:

$$T_{sig} = \bigcap_{g \in G\_sig} T_g$$

A spot is excluded from the signature footprint only if EVERY signature gene is a true zero there. Strict AND-gating would correspond to the union $\bigcup_g T_g$, which removes a spot as soon as any single gene drops out — creating the Swiss-cheese pattern that motivated this design. The intersection rule is strictly weaker: it requires unanimous agreement across the signature that a location is genuinely outside the co-expression territory. At $K = 2$ this reduces to "keep the spot unless both genes are true zero there", which is a sensible behavior on sparse Visium data with high dropout rates.

#### S6.3 Layer 2 — pixel-space mask after interpolation

Gaussian diffusion in EBIDW (S4.2) can leak non-zero signals from surrounding positive spots into true-zero regions. To block this *post-hoc*, NaVis projects $T_{sig}$ into pixel space, dilates it by $2 \cdot \sigma_{eff}$ pixels to cover the diffusion neighborhood, and zeros the per-gene maps at masked pixels before combining. Formally, let $P_{sig} = \text{dilate}(\text{project}(T_{sig}), 2\sigma_{eff})$.

$$E_g(p) \leftarrow 0 \text{ if } p \in P_{sig}$$

#### S6.4 Layer 3 — pixel union mask on combined signature

The combined signature is the arithmetic mean of the per-gene maps, each individually normalized to [0, 1]:

$$E_{sig} = (1 / K) \sum_{g \in G\_sig} \text{normalize}(E_g)$$

A final pixel-level union mask keeps any pixel where at least one per-gene map exceeds a small threshold ε = 0.01:

$$E_{sig}(p) \leftarrow 0 \text{ if } \forall g : E_g(p) \leq \varepsilon$$

This catches residual diffusion bleed-through that survived layers 1 and 2, while keeping the signature at any pixel where genuine expression remains.

### S7. Insights

Given any active channel $E$ (a gene map, signature, or H&E-blended overlay) and the three tissue priors, the Insights panel computes five compact readouts, each with an explicit biological question as its motivation.

#### S7.1 Compartment enrichment

Is the channel over- or under-represented in each compartment? For compartment $c$ with binary mask $M_c$ (density ≥ median(density[density>0])), the enrichment ratio is

$$r_c = \text{gmean}(E \mid M_c = 1) / \text{gmean}(E \mid \text{expressing})$$

where gmean is the geometric mean over expressing pixels only (pixels where $E > 0$). Expressing-only filtering ensures the ratio is not dominated by zero-expression pixels, which make the denominator near zero for sparse genes. The null distribution is obtained by 200 permutations of the compartment mask over expressing pixels:

$$p_c = (1 + |\{r_c^{(\pi)} : |r_c^{(\pi)} - 1| \geq |r_c - 1|\}|) / (1 + 200)$$

with the Laplace pseudocount preventing reported $p = 0$.

#### S7.2 Expression boundaries

Do sharp transitions in expression coincide with tissue edges? We extract the gradient magnitude of $E$ using the 3×3 Sobel operator:

$$\|\nabla E\| = \sqrt{(K_x * E)^2 + (K_y * E)^2}$$

with Sobel kernels $K_x$ = [[−1,0,1], [−2,0,2], [−1,0,1]] and $K_y = K_x^T$. We report the Pearson correlation of $\|\nabla E\|$ with each compartment's gradient magnitude $\|\nabla M_c\|$. High positive $r$ means expression boundaries track the compartment's structural transitions; $r$ near 0 means the gene is spatially organized at a scale independent of that compartment's edges.

#### S7.3 Spatial cross-correlation at lag distances

At what physical distance does the channel's pattern peak against each tissue prior? For a shift h in physical units (µm), the lag-h cross-correlation is

$$\rho_c(h) = \mathrm{corr}(E(p), M_c(p + h))$$

computed over all pixels where both operands are defined after the shift. The shift h is applied in the axis that maximizes the correlation (horizontal, vertical, or diagonal), reported as the signed scalar lag. We evaluate at lags 0, 25, 50, 100, 200, 400 and 800 µm, which ensures sufficient cover of the resolution of Visium (55 µm spot diameter) through biological distances. Lag 0 reports local co-localization; non-zero lags report whether the gene's pattern is offset from, or anti-correlated with, the compartment at that distance.

#### S7.4 Expression islands

Where are the high-expression foci, how many are there, how large, how round? We threshold $E$ at its 75th percentile over expressing pixels and extract connected components via 4-connectivity flood fill (`EBImage::bwlabel`). For each component $C_k$ of area $A_k$ (pixels) and perimeter $P_k$ (pixels) we report

$$\text{area}_{\mu m^2} = A_k \cdot (\mu m/\text{pixel})^2$$

$$\text{circularity} = 4\pi \cdot A_k / P_k^2$$

µm/pixel is derived from the Visium scale factor. Circularity is 1 for a perfect disc and approaches 0 for elongated or irregular components. We report total count, median and mean of each metric, and the min/max islands' areas.

#### S7.5 Histology decoupling score

To what extent is the channel's spatial pattern NOT explained by tissue architecture? We fit the linear model

$$E(p) = \beta_0 + \beta_1 M_{nuc}(p) + \beta_2 M_{fib}(p) + \beta_3 M_{soft}(p) + \varepsilon(p)$$

over pixels where $E > 0$ (expressing pixels only — decoupling is a property of where the gene exists, not of empty tissue). Let $R^2$ be the coefficient of determination. The decoupling score is

$$D = 1 - R^2$$

with interpretation bands: $D < 0.40$ = LOW (architecture largely explains the pattern), $0.40 \leq D < 0.80$ = MODERATE (partly decoupled), $D \geq 0.80$ = HIGH (pattern largely independent of architecture — a candidate for biology irreducible to H&E). These cutoffs are interpretability aids, not statistical claims.

### S8. Transcriptome-wide spatial similarity search

Given a query gene $q$, we find the $G-1$ genes whose full spot-level expression vector is most spatially similar to the query's. The critical design choice is to compute Pearson correlation over the entire spot set including zero-expressing spots:

$$\rho(q, g) = \mathrm{corr}(X_{q,\cdot}, X_{g,\cdot})$$

where the correlation is evaluated over all $N$ spots. The all-spot formulation gives high correlation only to genes whose expression pattern — ON and OFF together — matches the query's, which is the correct semantics for a spatial similarity search.

#### S8.1 BLAS-accelerated batch computation

Center $X$ row-wise (subtract per-gene mean across spots) and scale each row to unit $L_2$ norm; call the result $\bar{X}$. Then the full correlation vector between query $q$ and every other gene is a single matrix–vector product:

$$\rho = \bar{X} \cdot \bar{x}_q^{T}$$

which is implemented by a single call to BLAS *dgemv* in $O(GN)$ flops. On a typical Visium section with $G \approx$ 18 000 and $N \approx$ 4 000 this completes in under 5 seconds on a laptop CPU, with runtime dominated by the prep step (row-centering), not the matmul. The top 10 correlations are selected by partial sort.

### S9. Spot analysis

A click at pixel $p_0$ opens a drawer reporting five synchronous readouts at that location:

#### S9.1 Local H&E crop and color histogram

A 64×64 pixel crop of the H&E centered on $p_0$ is extracted and displayed. The RGB intensity distribution over the crop is shown as overlaid density estimates — a quick check of histological identity (purple-dominant = nuclei, pink-dominant = stroma).

#### S9.2 Tissue composition

The three tissue priors have different intrinsic scales: $M_{nuc}$ is a label image (integer connected-component IDs), whereas $M_{fib}$ and $M_{soft}$ are continuous densities in [0, 1]. To report comparable compositional fractions at a clicked spot, NaVis first binarizes the nuclei prior (any non-zero label = nuclei present), then takes the mean of each prior over a 4% radius neighborhood, and normalizes the three means to sum to 100%:

$$\rho_c(p_0) = \mathrm{mean}_{p \in \mathrm{patch}}\, \tilde{M}_c(p)$$

$$\mathrm{frac}_c = \rho_c \,/\, \textstyle\sum_{c'} \rho_{c'}$$

where $\tilde{M}_{nuc}$ is the binarized nuclei mask.

#### S9.3 Gene expression at spot

The nearest readout location to $p_0$ is identified by Euclidean distance on the pixel-coordinate map. Its raw count and CPM-normalized expression for every gene in the active channel are tabulated.

#### S9.4 Per-compartment channel correlation

For reference, the Pearson correlation of the entire active channel image with each compartment prior (computed over all expressing pixels) is displayed as a bar plot. This complements the lag-0 entry of the *Insights* cross-correlation readout and allows the user to confirm whether the slected location reflects the global channel–compartment relationship.

### S10. Default parameter table

All parameters exposed to the user through sliders are listed here, with defaults and sensible ranges. Parameters not listed are hard-coded constants.

| Parameter | Default value | Range |
|---|---|---|
| $\sigma_0$ (EBIDW base) | 10 | 1 –20 |
| n_iter (EBIDW) | 3 | 1 – 10 |
| t (sigmoid threshold) | 0.5 (manual) / 0.2 (ML) | 0 – 1 |
| sharpness | 30 | 1 - 50 |
| $\alpha$ (leakage) | 0.1 | 0 – 0.3 |
| $\tau$ (dropout – only effective for signatures) | 0.5 | 0 – 1 |
